# Paracoccin overexpression in *Paracoccidioides brasiliensis* reveals the influence of chitin hydrolysis on fungal virulence and host immune response

**DOI:** 10.1101/515056

**Authors:** Relber Aguiar Gonçales, Vanessa Cristina Silva Vieira, Rafael Ricci-Azevedo, Fabrício Freitas Fernandes, Sandra Maria de Oliveira Thomaz, Agostinho Carvalho, Patrícia Edivânia Vendruscolo, Cristina Cunha, Maria Cristina Roque-Barreira, Fernando Rodrigues

## Abstract

*Paracoccidioides brasiliensis* and *P. lutzii*, etiological agents of paracoccidioidomycosis (PCM), develop as mycelia at 25-30 °C and as yeast at 35-37 °C. Only a few *Paracoccidioides* spp. proteins are well characterized. Thus, we studied paracoccin (PCN) from *P. brasiliensis*, its role in the fungus biology, and its relationship with the host innate immune cells. Cloning and heterologous expression analysis revealed its lectin, enzymatic, and immunomodulatory properties. Recently, we employed a system based on *Agrobacterium tumefaciens*-mediated transformation to manipulate *P. brasiliensis* yeast genes to obtain clones knocked-down for PCN, which after all, are unable to transit from yeast to mycelium forms, causing a mild pulmonary disease. Herein, we generate *P. brasiliensis* overexpressing PCN (ov-PCN). To date, it was not explored the overexpressing of endogenous components in *Paracoccidioides* spp. Therefore, we investigate the role of PCN in fungal biology and pathogenesis. Augmented levels of PCN mRNA and protein, and N-acetylglucosaminidase activity confirmed PCN overexpression in ov-PCN of *P. brasiliensis* yeasts. Interestingly, PCN overexpression did not affect the yeasts’ growth or viability and favored cell separation. The ov-PCN clones transitioned faster to the mycelium form than the wt-PCN yeasts. Concerning infection, while most of mice infected with the wt-yeasts (90%) survive at least until the 70^th^ day, all mice infected with ov-PCN yeasts were already died at the 35^th^ day post-infection. In vitro assays showed that ov-PCN were more susceptible to phagocytosis by macrophages. Finally, it was verified that the chitin particles isolated from the ov-PCN cells were smaller than those obtained from the wt-PCN yeasts. Macrophages stimulated with the chitin isolated from ov-PCN produce IL-10, whereas the particles with a wider size range harvested from wt-PCN yeasts induced TNF-α and IL-1β secretion. The anti-inflammatory microenvironment from macrophage stimulation with small chitin particles hampers the development of a protective immune response against the fungus. We postulated that the high grade of chitin cleavage, as the results of augmented PCN expression, favors pathogenesis following *P. brasiliensis* infection. Thus, PCN is a relevant virulence fungal factor.

**AUTHOR SUMMARY:** *Paracoccidioides* spp. are pathogenic fungi that cause paracoccidioidomycosis (PCM) in humans, the main deep mycosis of Latin America. Recently, by knocking down the paracoccin gene, our group showed that this lectin is necessary for the morphological transition from yeast to hyphae, and that this decrease results in low *P. brasiliensis* virulence. Here, after overexpress PCN, we revealed the importance of the yeast chitin hydrolysis to the host response. Infection of mice with ov-PCN yeasts causes severe lung disease compared to moderate disease caused by wt-PCN yeasts. The release of smaller chitin particles was as a result of an accelerated chitin hydrolysis provided by ov-PCN yeasts. Interestingly, these smallest chitin particles are able to modulate host response by increasing IL-10 in the meantime that decrease TNF-α secretion, thus hampering Th1 immune response that is crucial in the fight against this fungi. These findings represent a significant advance in the knowledge about the role of PCN chitinase in *P. brasiliensis*.

## INTRODUCTION

Paracoccidioidomycosis (PCM) is a severe mycosis widespread in Latin America [1]. The fungi that cause PCM belong to the genus *Paracoccidioides*, which include two species of thermo-dimorphic fungi developing as filaments at 25-30 °C, mainly in the soil, and assuming the yeast form at 35-37 °C [2]. *Paracoccidioides lutzii* is endemic in the Midwest and Southeast regions of Brazil, whereas *Paracoccidioides brasiliensis* is found in Southeast and North Brazil [3-5].

The virulence mechanisms of *Paracoccidioides* spp. are poorly known owing to the lack of adequate characterization of fungal components. Among them, we highlight the glycoprotein gp43 [6-10], Hsp60 [11-13], Pb40 [14], Pb27 [15], SconCp [16], Cdc42p [17] and paracoccin (PCN) [18-20]. We described PCN from *P. brasiliensis* as a major protein component of a GlcNAc-binding fraction of yeast extracts [19, 20]. The preparations obtained by affinity to immobilized GlcNAc or chitin allowed us to identify several properties of PCN: (*i*) it contributes to fungal adhesion to extracellular matrix (ECM) components, such as laminin, in a carbohydrate recognition- and concentration-dependent manner [19]; (ii) it induces macrophages to produce TNF-α and nitric oxide [19]; (iii) it exerts N-acetylglucosaminidase (NAGase) activity, which accounts for yeast growth and cell wall biogenesis [21, 22].

The only established molecular system of *Paracoccidioides* spp. genetic manipulation is based on *Agrobacterium tumefaciens*-mediated transformation (ATMT). It was adopted by the few research groups working on transformation of these fungi [16, 17, 23-29]. The method uses T-DNA binary vectors harboring user-defined genetic constructs, which have already allowed the expression of an exogenous marker, the green fluorescent protein [23] or to down-regulate gene expression by antisense RNA (*aRNA*) [25]. The publication of the complete genome sequence of three isolates (Pb18, Pb03, and Pb01) from the *Paracoccidioides* genus [30, 31] allowed further knowledge on genome of these species opening new targets for study. In recent studies [28, 29], strains with downregulation of Pb14-3-3 and PbSOD3 genes were developed by using *aRNA* through ATMT technology. It was shown that the Pb14-3-3 contributes for the fungal attachment to ECM/pneumocytes and fungal virulence [28], whereas PbSOD3 is essential for events underlying host-pathogen interaction [29].

Recently, our group successfully knocked-down PCN expression. Working with the obtained *P. brasiliensis* transformed yeasts, we verified that PCN is essential for the fungal morphological transition and virulence [18]. To date, overexpression of endogenous components has not been explored in *Paracoccidioides* spp. In this work, we induced the overexpression of PCN in *P. brasiliensis* yeasts, which were used to investigate the role of PCN in fungal biology and the fungus-host interaction. We unravel the mechanism accounting for the severe pulmonary disease caused by yeasts overexpression PCN in mice, through the differences in the structure size of chitin.

## RESULTS

### Increased PCN expression by co-culturing *P. brasiliensis* yeasts with macrophages

In a recent study, we compared the PCN transcripts in different morphotypes of *P. brasiliensis.* Although detected in all fungal forms, PCN mRNA expression was high in hypha and yeast-to-hypha transition forms [32]. Herein, we investigated PCN expression in yeasts for 24 h during THP-1 human macrophages infection. The PCN expression was 9,000-fold increased upon infection (Fig 1A). In addition, as evaluated by western blotting, PCN protein content was detectable in the supernatant of yeast cultures and well visualized in the supernatant of yeast/macrophage co-cultures (Fig 1B). These results prompted us to investigate the implications of PCN overexpression in fungal biology and host immunity.

**Fig 1.**
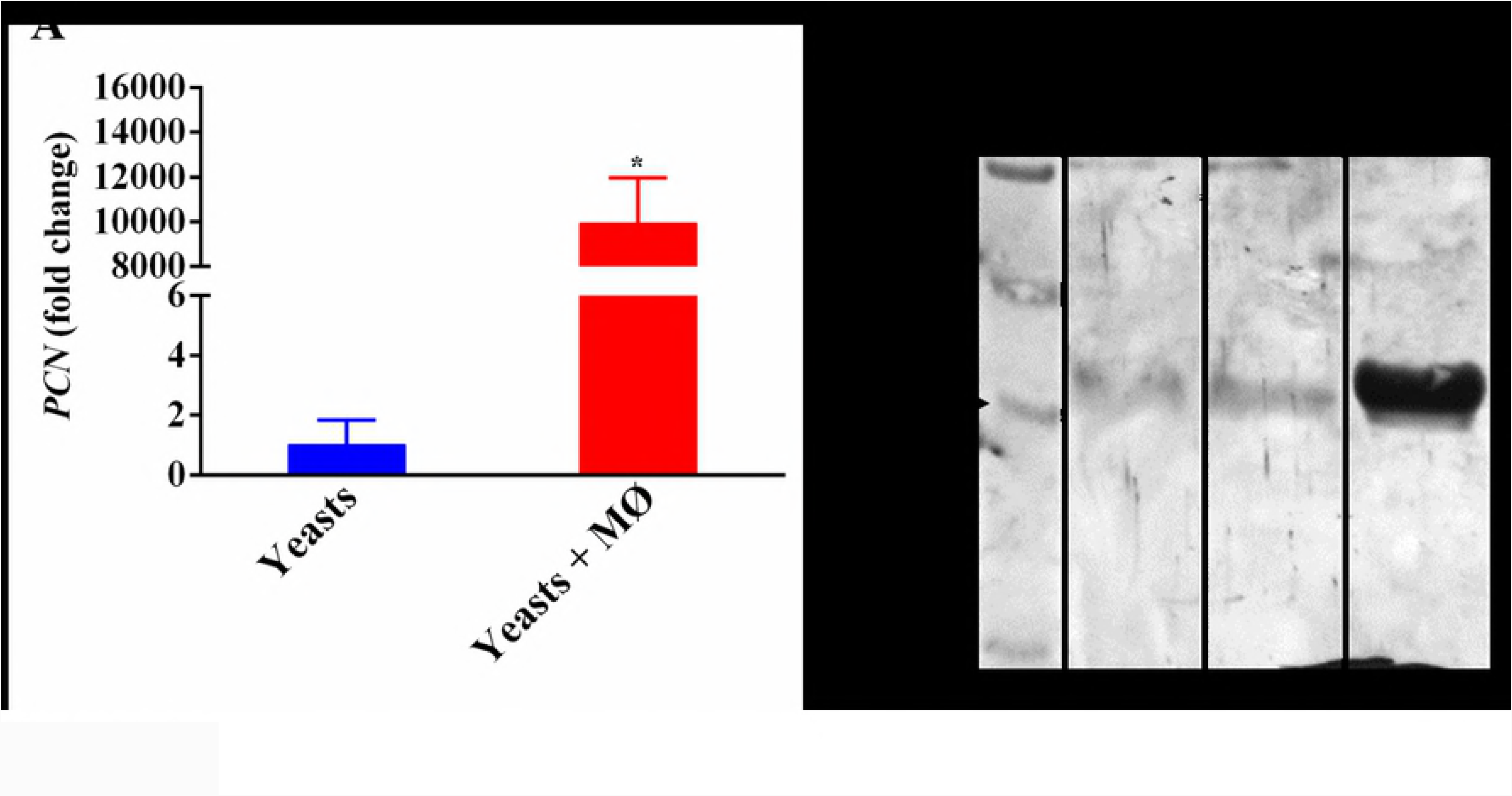
*P. brasiliensis* yeasts co-cultured with human macrophages augment PCN expression. **(A)** Augmented PCN expression in *P. brasiliensis* yeasts co-cultured for 24 h with THP1 human macrophages. The levels of PCN mRNA expression were determined by qRT-PCR and normalized to the expression level of the internal references TUB2 and L34. Bars depict the mean ± SEM fold change in PCN expression found in co-cultured yeasts with macrophages in relation to the PCN expression verified in yeasts cultured alone. The comparison was done through the One-Way ANOVA test. **\****p* < 0.1. **(B)** Western blot analysis of a recombinant PCN protein, expressed in *Pichia pastoris* [48] and the PCN content in the supernatant of yeast cells that were co-cultured or not with THP-1 macrophages. Reactions were developed by IgY anti-PCN conjugated to HRPO. MW, molecular weight.

### Characterization of PCN-overexpressing yeasts

PCN is structurally related mostly to chitinases from *Histoplasma capsulatum* (83% similarity), *Blastomyces dermatitidis* (81% similarity), and class III ChiA1 of *A. nidulans* (83% similarity) (*NCBI – National Center for Biotechnology Information*). To generate PCN-overexpressing *P. brasiliensis* yeasts, we used the ATMT methodology, after cloning the genomic sequence of a hypothetical sequence PADG_03347 (gPCN) under the RHO2 promoter control in transfer DNA (T-DNA). The yeast cells of the Pb18 wild type strain were transformed with T-DNA containing gPCN (Fig 2A) or empty vector. We confirmed the integration of the T-DNA cassette as well as the mitotic stability in six randomly selected transformants (gPCN; data not shown). We demonstrated, by comparing PCN mRNA levels determined by qRT-PCR, that the isolated clones expressed a range from 1.5-to 7-fold more PCN than the wt-PCN yeasts (Fig 2B). The isolated clones were re-named according to the fold increase in PCN expression (gPCN7 > gPCN6 > gPCN4). For those clones expressing higher levels of PCN we determined the specific activity of the N-Acetyl-β-D-glucosaminidase [20] in the supernatant of the yeast cultures. As shown in Fig 2C, the supernatants of the cultures of the gPCN yeasts provided significantly higher N-Acetyl-β-D-glucosaminidase activity than those detected in the supernatant of wt control yeasts. Once proven the overexpression of PCN in the selected clones, their growth, viability, and size of yeasts were determined.

**Fig 2.**
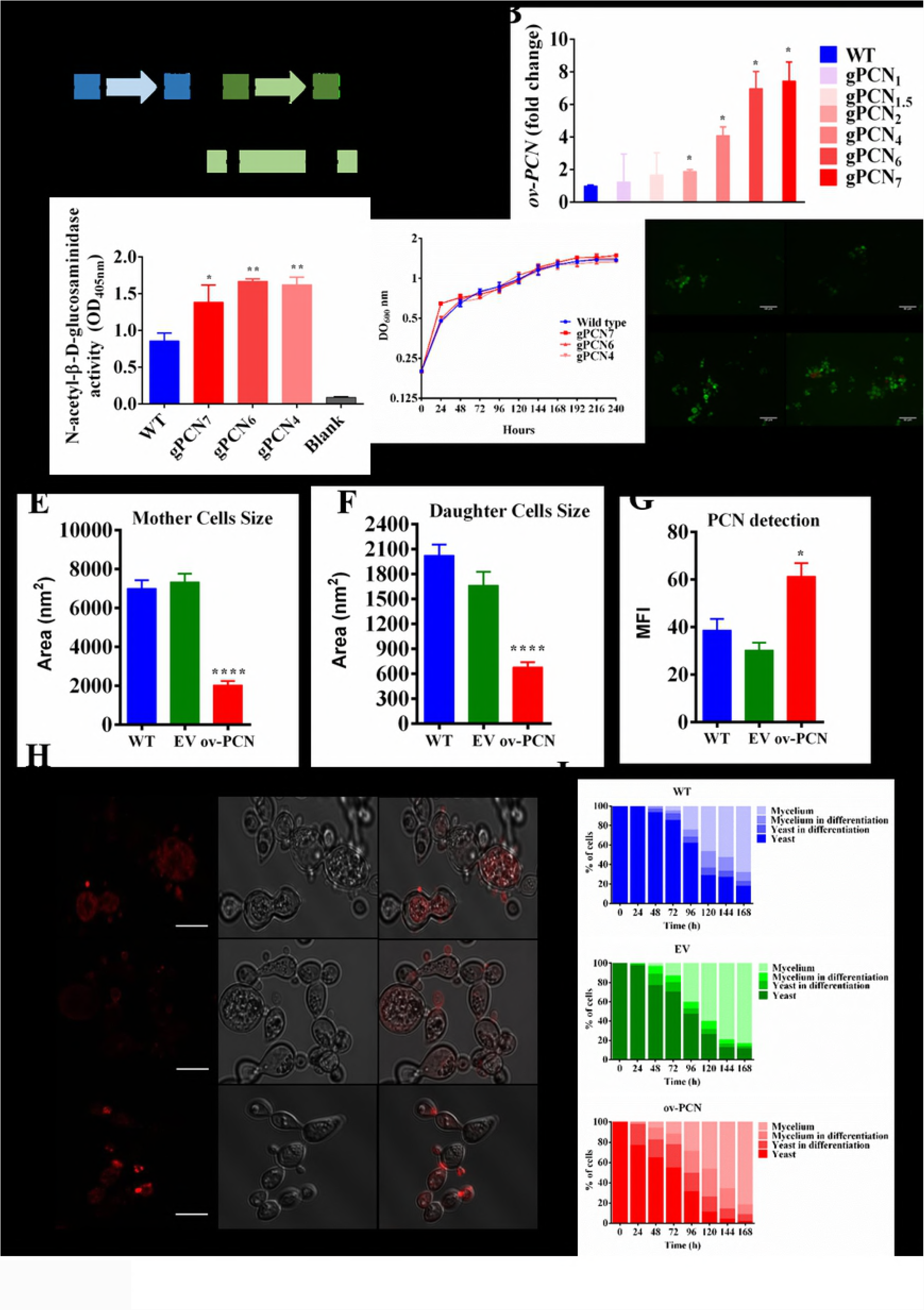
Characterization of ov-PCN yeasts generated by using the pUR5750 plasmid and a transformation system mediated by *Agrobacterium tumefaciens*. **(A)** The PCN overexpression cassette. The genomic Paracoccin sequence (PADG_03347) was produced based on the Pb339 (PbWT) genomic sequence, as detailed in the Materials and Methods section, and cloned under the control of the Rho2 ATPase promoter from *P. brasiliensis* (RHO2). The constructs were sub-cloned into the T-DNA region of the binary vector pUR5750 harboring the *E. coli* hygromycin B phosphotransferase (HPH) resistance gene driven by the glyceraldehyde 3-phosphate promoter from *A. nidulans* (PGPDA) and bearing transcriptional terminators (T) from cat-B and TRPC; LB: left border; RB: right border. **(B)** Gene expression levels of gPCN yeasts and controls wt-PCN and EV after subculture for 3 days (gene expression levels obtained by qRT-PCR were normalized to the level of expression of the internal reference, TUB2, and L34); **\****p* < 0.01 compared with wt-PCN strains. **(C)** Supernatants of gPCN yeasts and wt-PCN strains, cultured in DMEM, were assessed for NAGase activity, detected by the degradation of the *p*-nitrophenyl-N-acetyl-β-D-glucosaminide substrate. **(D)** gPCN yeasts and wt-PCN controls had their growing evaluated by culture turbidity, as determined by the OD_600_ reading; they were also examined for yeast cell viability by diacetate fluorescein and ethidium bromide [53]. **(E and F)** The area (in nm^2^) of each mother and daughter yeast cells harvested on the third day of culture (exponential growth) was determined by using the ImageJ software (http://rsb.info.nih.gov./iji/). A total of 50 cells were examined per slide, by confocal microscopy. **(G)** Fluorescence intensity of yeasts as detected by confocal microscopy and analyzed with the aid of the ImageJ software. **(H)** The ov-PCN yeasts and the respective control yeasts (wt-PCN and EV) grew in solid BHI medium supplemented with 4% FBS for 72 h (exponential phase). The images were obtained in a confocal fluorescence microscope after the samples were incubated with IgY anti-PCN conjugated to Alexa Fluor 594 fluorophore. Bars correspond to 200 µm. **(I)** Frequency of transition forms for either transformants or control yeasts was determined as previously reported [16, 58] through quantitative optical microscopy (40 × magnification). Results were expressed as the percent of total counted cells (n=300). Graphical representation of the morphological signs of (Y→M) transition occurred during the 7-days culture period at 25 °C. The assay was performed in triplicate and previous established patterns were utilized to drive the fungal forms classification [16, 58]. Bars depict the mean ± SEM. Results provided by the ov-PCN and wt-PCN yeasts were compared through the One-Way ANOVA test. \*\*\*\**p*<0.0001, \*\*\**p*<0.001, \*\**p* <0.01 and \**p*<0.1.

To compare the growth profile of the gPCN and wt-PCN yeasts, the cellular density of cultures in BHI was evaluated along 10 days by regular readings of the optical density at OD_600nm_. Our data shows overlapped time-course curves for all gPCN and wt-PCN yeasts, indicating that PCN overexpression did not affect fungal growth. We also examined cell viability by fluorescence microscopy, after staining with diacetate fluorescein and ethidium bromide. Our data shows no differences in viability (Fig 2D). Because we verified no significant difference in the three selected gPCN regarding their general characteristics, we selected gPCN7 for further experiments and it is hereafter identified simply as ov-PCN yeast. The measurement of the cell-area (cell size) of mother and daughter yeast cells showed that ov-PCN yeasts had a smaller size than the wt or empty vector (EV)-transformant yeasts (Fig 2E and F).

PCN was quantified in ov-PCN and wt-PCN yeasts by confocal microscopy. After incubation with IgY anti-PCN conjugated to Alexa Fluor 594, PCN was quantified in the yeast samples by measuring the fluorescence intensity, with the aid of the ImageJ software. Accordingly with the data obtained for the the N-Acetyl-β-D-glucosaminidase activity (Fig 2C), fluorescence was higher in the ov-PCN yeasts, compared to wt-PCN and EV control yeasts (Fig 2G). The yeast samples were also analyzed for cell wall PCN distribution. In both, ov-PCN and wt-PCN yeasts, PCN was detected along the yeast cell wall, and primarily concentrated in the budding regions (Fig 2H). These fungal structures were more intensely fluorescent in ov-PCN than in wt-PCN yeasts.

Recently, we have shown that PCN-silenced yeasts of *P. brasiliensis* are unable to perform the morphological transition from yeast to mycelium [18]. Herein, we studied the transition of ov-PCN yeasts by quantitative optical microscopy, for 7 days (168 h) following the shift in culture temperature from 37 to 25 °C. We observed that, at 24 h, more than 20% of the ov-PCN yeasts showed morphological signs of differentiation, which increased to 50% at 72 h. The transition to mycelium in 90% of the ov-PCN yeasts completed during the experimental period of 7 days. Meanwhile, the control cultures of wt-PCN yeasts or EV-transformed yeasts required 96 to 120 h to have 50% of fungal cells with morphological signs of differentiation; furthermore, during the total experimental period of one-week, differentiation had not even started for 10 to 20% of wt-PCN yeasts (Fig 2I). These results show a faster ov-PCN yeast transition compared to wt-PCN or EV. Our data supports the critical effect of PCN in the morphological transition from yeast to mycelium and cell wall biogenesis of *P. brasiliensis* [21, 22, 32].

### Pathogenic features of the PCN-overexpressing yeasts in mice

Because the *P. brasiliensis* PCN-knocked down yeasts are less virulent than the wt-PCN yeasts [18], we hypothesized that overexpression of PCN could promote different pathogenic profile of yeast in mice. The lungs of BALB/c mice, inoculated through the intranasal route with 2×10^6^ cells of ov-PCN or wt-PCN yeasts, were microscopically examined, thirty days after infection, regarding the extension of the granulomatous lesions. In mice that were infected with wt-PCN yeasts, small and circumscribed granulomas were focally distributed in the pulmonary tissue, whereas in mice infected with ov-PCN yeasts, the granulomas were large and coalescent, occupying an extended area of the lungs (Fig 3A). Methenamine/silver-staining of the pulmonary sections revealed few and focally distributed viable yeasts, more centrally localized in the small and compact granuloma of wt-PCN yeast-infected mice, while abundant viable yeasts were dispersed in all the area of coalescent granulomatous lesions of the mice infected with ov-PCN yeasts (Fig 3B). Morphometric analysis of pulmonary tissue injury showed that the area occupied by lesions was 60% larger in mice that were infected with ov-PCN yeasts than in wt-PCN yeast-infected mice (Fig 3C). Pulmonary CFU counting has quantitatively validated the results obtained by optical microscopy; the number of colonies provided by the ov-PCN yeast-infected mice was at least one order of magnitude higher than the one obtained from wt-PCN yeast-infected mice (Fig 3D). In our first analysis of the mechanisms accounting for the different profile of infection we considered that the ability of *P. brasiliensis* to cause disease largely depends on the yeasts’ resistance to the defense mechanisms of the host phagocytes [33].

**Fig 3.**
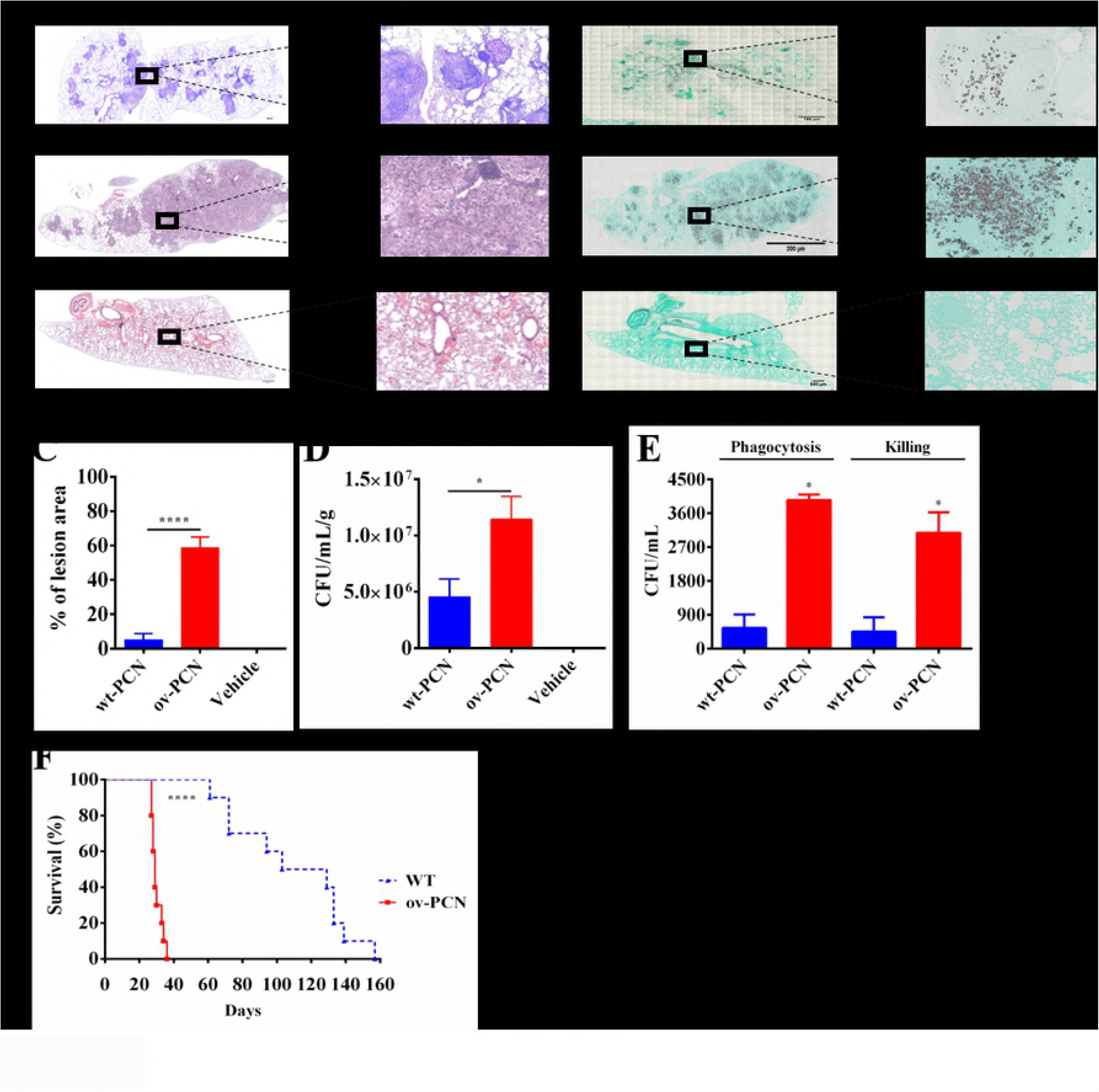
Pulmonary fungal burden, granulomatous lesions, and mortality of infected mice with ov-PCN *P. brasiliensis* yeasts. BALB/c mice were infected by intranasal route with 2×10^6^ ov-PCN or wt-PCN *P. brasiliensis* yeasts. On day 30 after infection, a pulmonary lobe of each animal was assessed for the recovery of pulmonary CFU. **(A)** Each group was consisted of five mice. A non-infected control group (mice inoculated with a vehicle, PBS) was included. The sections were stained with hematoxylin and eosin or **(B)** with Gomori’s methenamine silver to visualize the viable yeasts. Images were captured using a Carl Zeiss Axiophot microscope coupled to a JVC TK1270 camera. Magnification bars: 500 nm for images of the whole organ sections and 200 nm for partial images of lung sections. **(C)** The area (%) of lung sections occupied by granulomatous lesions, was quantitated with the aid of the ImageJ software. **(D)** CFU recovery from the lung homogenates of infected mice with ov-PCN or wt-PCN yeasts. **(E)** Murine macrophages of the RAW 264.7 cell lineage (1×10^6^ cells/mL) were infected with ov-PCN or wt-PCN *P. brasiliensis* yeasts (MOI=1:10). At 4 and 48 h post-infection, macrophages were washed and lysed. The lysates were assessed for CFU recovering, whose results allowed to estimate the yeasts’ sensitivity to the macrophages effector functions, phagocytosis (4 h post-infection) and killing (48 h post-infection). The results are expressed as mean ±SEM and compared between ov-PCN and wt-PCN yeasts through two-tailed t-test analysis of variance. **(F)** Survival curve of male BALB/c mice infected with ov-PCN or wt-PCN yeasts (1×10^7^ yeasts per animal) that were monitored daily for death occurrence over a 160 day post-infection period. Mice survival are expressed in percentage. The assays were carried out in triplicates. Bars depict the mean ± SEM and the values provided by infected mice with ov-PCN yeasts were compared to values obtained from wt-PCN yeast-infected mice by the One-Way ANOVA test. \*\*\*\**p* < 0.0001 and \**p* < 0.1.

As such, we assayed *in vitro* the sensitivity of ov-PCN and wt-PCN yeasts to RAW 264.7 murine macrophage effector functions. Phagocytosis was examined 4 h after incubating macrophages with ov-PCN or wt-PCN yeasts. Our data indicate that phagocytosis of ov-PCN yeasts is more effective than that of wt-PCN yeasts (Fig 3E). The fungal killing by macrophages was investigated by obtaining the lysate of macrophages that were incubated for 48 h with yeasts. We recovered a significantly higher CFU number from the cells infected with ov-PCN yeasts than with wt-PCN yeasts because a higher number of yeasts were internalized by macrophages infected with ov-PCN yeasts than wt-PCN (Fig 3E).

Concerning the mice survival to the fungal infection, we observed that all animals infected with wt-PCN yeasts survived throughout a 40 days post-infection period, whereas the ones infected with ov-PCN yeasts started dying at day 27 post-infection and none survived to a 35 days post-infection period (Fig 3F). We concluded that PCN overexpression aggravates the disease caused by *P. brasiliensis* yeasts, reinforcing the notion that PCN acts as a *P. brasiliensis* virulence factor [18].

### Effect of PCN overexpression on the chitin content of the yeast cell walls

Having defined the augmented levels of the PCN protein in ov-PCN yeasts (Fig 2) and considering the already known PCN properties of binding to chitin and exerting chitinase activity [20], we evaluated the chitin content on the cell wall of the strains under study. The staining of ov-PCN, wt-PCN, and EV yeasts with calcofluor white allowed detecting chitin by fluorescent confocal microscopy (Fig 4A). With the aid of the ImageJ software, we could verify that chitin detection was reduced by 40% in ov-PCN yeasts, in comparison to wt-PCN yeasts (Fig 4B). We also detected the chitin content of yeast cells by using a TexasRed conjugate to Wheat Germ Agglutinin (WGA), a highly specific chitin-binding lectin [34]. The analysis by flow cytometry also showed a significant reduction (by 30%) of the chitin content in ov-PCN yeasts, compared to that detected in wt-PCN yeasts (Fig 4C). Consistently the cell wall of wt-PCN yeasts, as examined by electron microscopy, was about 6-fold thicker than that of ov-PCN yeasts (Fig 4D and 4E).

**Fig 4.**
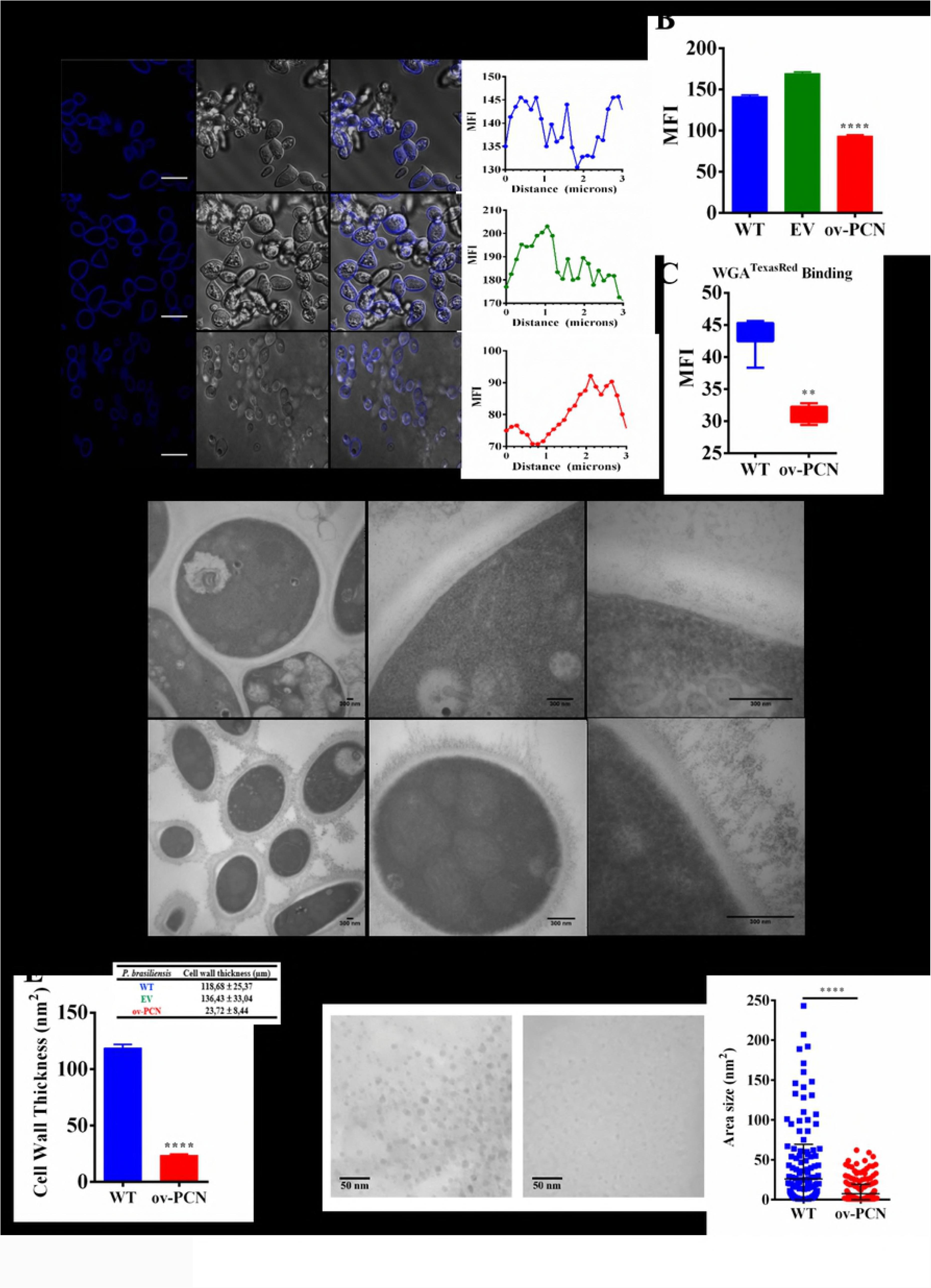
PCN overexpression in *P. brasiliensis* yeasts correlates with diminished chitin content, narrowed cell wall, and detection of small chitin particles in the supernatant of yeast cultures. **(A)** ov-PCN yeasts and the controls wt-PCN and EV yeasts, grown in solid BHI broth for 72 h (exponential phase) and stained with calcofluor white (10 mg/mL) were assessed for their chitin content in the cell wall by fluorescence confocal microscopy. Bars correspond to 200 µm. The mean fluorescence intensity (MFI) and the distances (micron) of the fluorescence pics from the nucleus were determined with the aid of the ImageJ software and graphically represented (at the right side of panel A). **(B)** The MFI of the yeasts was determined with the aid of the ImageJ software. **(C)** To assess the amount of chitin present in the yeast cell wall, 500 µL samples of culture supernatants of ov-PCN or wt-PCN yeasts were harvested at 72 h (exponential growth). The samples were incubated for 30 min with WGA conjugated with a TexasRed fluorophore. Analysis was performed by flow cytometry. **(D)** Electron microscopy of the cell wall of wt-PCN and ov-PCN yeasts. Bars correspond to 300 nm. **(E)** The cell wall area (nm^2^) was analyzed with the aid of the ImageJ software. The results provided by the EV and wt-PCN yeast cells were similar. **(F)** The diameter of chitin fragments isolated from the ov-PCN and wt-PCN cultures by using a column with WGA and the area size (nm^2^) and integrated density (pixels) was analyzed by the ImageJ software. Bars depict the mean ± SEM and were compared by Mann-Whitney’s test, \*\*\*\**p* < 0.0001, and ** *p* <0.0022.

Then, the size of the chitin particles present in the supernatant of yeast cultures and captured by immobilized WGA was analyzed by electron microscopy. The isolated chitin particles had variable sizes (Fig 4F). We detected only particles with less than 60 nm^2^ in the material derived from the supernatants of ov-PCN yeasts. This low size range was also prominent in the material obtained from the wt-PCN yeasts; nevertheless, this preparation included a wider size distribution, which included particles with areas as large as 240 nm^2^ (Fig 4F).

Finally, the obtained results consolidate the notion that PCN hydrolyzes chitin of the *P. brasiliensis* cell wall, reducing its chitin content and cell wall thickness. In addition, the process triggered by PCN promotes the release of very small chitin particles to the extracellular milieu.

### Macrophage activation by chitin particles from ov-PCN yeasts

It was previously reported that the size of chitin fragments correlates with the particles’ property of stimulating macrophages to produce inflammatory or anti-inflammatory cytokines [35-41]. Based on these studies, we examined whether the chitin fractions we captured from the supernatants of ov-PCN (containing only small chitin fragments) or wt-PCN yeasts (containing a large spectrum of small and larger chitin fragments) (Fig 4F) could result in distinct responses from murine bone marrow-derived macrophages (BMDMs). Isolated chitin particles of wt-PCN and ov-PCN yeasts were assayed for the ability of inducing cytokine production by BMDMs. Dose-response and time-course curves (supplementary data, S1A-F) were drawn for TNF-α, IL-1β, and IL-10 cytokines, whose production was tested toward five different chitin concentrations (from 2.5 to 100 µg/mL) during the periods of 24, 48, and 72 h. The dose-response curves for the levels of TNF-α, IL-1β and IL-10, representing inflammatory and anti-inflammatory cytokines, respectively, which shows that the two curves are inverted: lower chitin concentrations correlate with higher IL-10 levels, whereas higher chitin concentrations determine higher TNF-α production (Fig 5). The production of the pro-inflammatory cytokines TNF-α (Fig 5A) and IL-1β (Fig 5B), measured in the supernatant of BMDMs harvested 48 h after stimulation, was significantly higher when stimulated by chitin particles derived from wt-PCN than ov-PCN yeasts. On the other hand, the IL-10 levels measured in the supernatant of BMDMs (Fig 5C), also harvested 48 h after stimulation, were significantly higher when stimulated with chitin particles derived from ov-PCN than wt-PCN yeasts.

**Fig 5.**
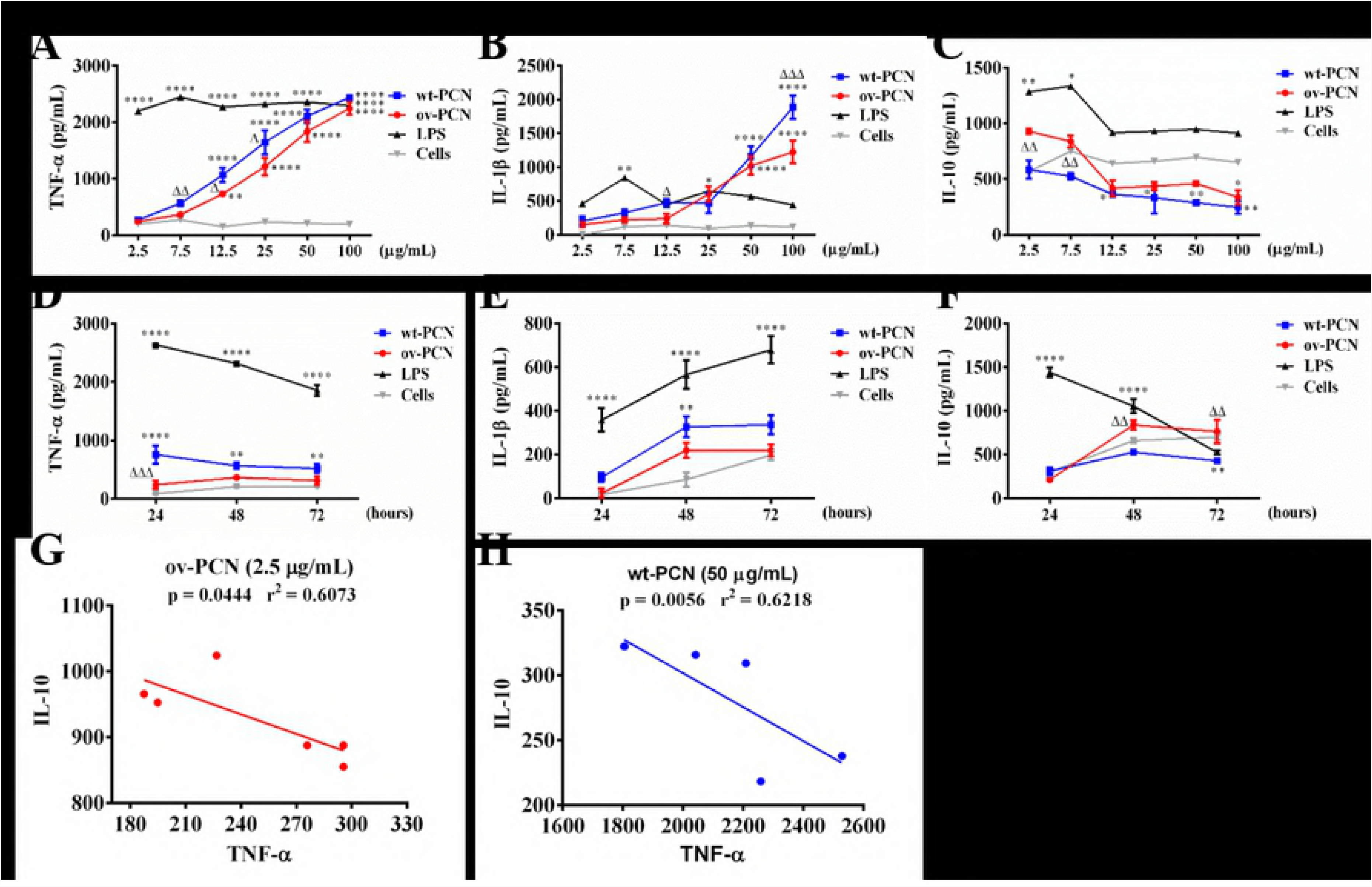
Chitin particles isolated from supernatants of ov-PCN and wt-PCN *P. brasiliensis* yeasts induce cytokine release. BMDMs (1x10^6^ cells/mL) were stimulated by chitin particles captured from the supernatant of ov-PCN or wt-PCN *P. brasiliensis* cultures. Following 48 h stimulation, the supernatants of BMDMs were assessed by ELISA for cytokines levels (pg/mL) in a dose-response **(A-C)** and time-response manner using 7.5 µg/mL of chitin particles **(D-F)**. **(A and D)** TNF-α. **(B and E)** IL-1β. **(C and F)** IL-10. Unstimulated and LPS-stimulated BMDMs were used as negative and positive controls, respectively. **(G and H)** Correlation between levels of IL-10 and TNF-α produced by BMDMs. The negative correlation between IL-10 and TNF-α levels in BMDMs stimulated with chitin particles isolated from the supernatant of ov-PCN (2.5 μg/mL) **(G)** or wt-PCN (50 μg/mL) **(H)**. Analyses were performed using the Spearman test, each point represents a sample, and the values of r and *p* are indicated in the graphs. The results are expressed as mean ± SEM and compared to the response to medium alone through one-way analysis of variance, followed by One-Way ANOVA test. The samples were compared to medium, \*\*\**p*<0.001, \*\**p*<0.01 and \**p*<0.1, and when we’re compared to ov-PCN with wt-PCN the values *p* are ^ΔΔΔ^*p*<0.001, ^ΔΔ^*p*<0.01 and ^Δ^*p*<0.1.

In order to demonstrate the regulation of cytokines mediated by the chitin fragments isolated from the supernatants of the ov-PCN and wt-PCN yeast cultures by BMDMs at different time points, we used the concentration of 7.5 μg/mL of chitin (Fig 5D-F). Cytokine induction by BMDMs exhibited increased levels of TNF-α and IL-1β from 24 h post-infection by chitin particles isolated from wt-PCN yeasts than macrophages stimulated with ov-PCN yeast-derived particles (Fig 5D and 5E). On the other hand, modulation by IL-10 was increased from 48 h and was maintained over time in BMDMs stimulated with chitins derived from ov-PCN yeasts; macrophages stimulated with wt-PCN yeast-derived particles had basal levels similar to the negative control (Fig 5F). When we performed correlation analyses using the data obtained in the dose-response experiment, we found that levels of IL-10 and TNF-α correlated negatively with concentrations of 2.5 μg/mL and 50 μg/mL of chitin particles isolated from the supernatant of ov-PCN and wt-PCN, respectively. From this, these results suggest that isolated particles of the ov-PCN supernatant (small chitins) induce higher levels of IL-10 and lower TNF-α than wt-PCN yeasts (Fig 5G), whereas particles isolated from the supernatant of wt-PCN yeasts (large chitins) induce lower levels of IL-10 and higher TNF than ov-PCN (Fig 5H). Taken together, these results demonstrate that all the chitin captured from the supernatant of cultures of *P. brasiliensis* wt-PCN yeasts, consisting of small and large particles, used at concentrations higher than 7.5 µg/mL, induce BMDMs to produce the pro-inflammatory cytokines TNF-α and IL-1β. Inversely, the chitin particles captured from the supernatant of ov-PCN yeasts cultures consisted exclusively of small chitin particles, used at concentrations as low as 2.5 to 7.5 µg/mL; these particles induced the basal production of the anti-inflammatory cytokine IL-10 in BMDMs. Our conclusion is consistent with literature data showing that small chitin particles induce IL-10 production by macrophages [35, 42], whereas larger chitin particles induce TNF-α release [35, 36, 38, 39, 42]. Because high levels of anti-inflammatory cytokines in the initial phase of *P. brasiliensis* infection induce the non-protective immune response [43], our findings explain, at least partially, the high severity of the pulmonary disease developed in mice that were infected with *P. brasiliensis* yeasts overexpressing the chitinase PCN.

## DISCUSSION

This is the first report of gene overexpression in a fungal species of the genus *Paracoccidioides*. The gene (annotated as PADG_03347) codes for PCN, whose previous cloning and heterologous expression allowed the identification of this multidomain protein that exerts biological activities of lectin and binds to GlcNAc and chitinase [19], which hydrolyzes chitin [20]. It also acts as an immunomodulatory agent [44, 45]. Yeast transformation was mediated by ATMT [23, 46], a system reported to be successful when employed to obtain *P. brasiliensis* knocked-down clones for proteins playing relevant roles in the fungal virulence or pathogenesis, e.g., CDC42 [17], PbHAD32 [24], asSconC [16], PbGP43 [26], PbP27 [27], Pb14-3-3 [28], and PbSOD3 [29]. By applying the ATMT methodology, we recently obtained PCN-knocked-down *P. brasiliensis* yeasts, which made possible identifying PCN as a fungal virulence factor [18]. Finally, in the present study, the ATMT system successfully provided PCN-overexpressing *P. brasiliensis* yeasts, that allowed us to confirm that PCN is a virulence factor that affects fungal pathogenesis and identify mechanisms accounting for the roles played by PCN. The ATMT methodology was also successful in terms of the mitotic stability of the generated ov-PCN yeasts.

Our interest on the PCN gene manipulation comes from demonstrations that the subcutaneous administration of recombinant PCN (rPCN) to infected mice with *P. brasiliensis* promotes modulation of the host immune response and confers protection against the fungal disease [44, 45]. The response is triggered by the PCN lectin domain interaction with N-glycans of TLR2 and TLR4 [45, 47]. Both receptors are expressed on the surface of macrophages, which undergo M1 polarization followed by high production of pro-inflammatory mediators, such as the Th1 polarizing cytokine IL-12 [48]. We demonstrated that the developed Th1 immune response accounts for the host resistance to the *P. brasiliensis* infection conferred by the PCN administration [45]. A subsequent study, focused on PCN distribution in *P. brasiliensis*, revealed its association with chitin and prominent localization in structures related to fungal growth, such as hyphae tips and budding regions of yeast cell wall [32]. The relevant biological activities and peculiar localization of PCN motivated us to investigate the role the endogenous PCN could play in fungal biology, as well as its effect on the host immune response.

The PCN overexpression strongly influences the fungal transition and the course of the murine *P. brasiliensis* infection. The overall data showing the opposite biological effects between PCN-overexpression and PCN-silenced fungi, [18], as shown in Table 4, further supports the need of a fine tuning PCN expression.

**Table 4.** General characteristics of PCN-silenced and ov-PCN *P. brasiliensis*.

| Features<br>Transformed yeast | Detection | Yeast Growth | Transition<br>(Y→M) | Killing by Mφ | Pathogenesis | Virulence |
| --- | --- | --- | --- | --- | --- | --- |
| PCN-silencing<br>(kd-PCN) | ↓ mRNA<br>↓ NAGase | Not affected | Blocked | Susceptible | ↓ Fungal burden<br>↓ Lung lesions | Reduced |
| PCN-overexpressing<br>(ov-PCN) | ↑ mRNA<br>↑ NAGase | Not affected | Accelerated | Susceptible | ↑ Fungal burden<br>↑ Lung lesions | Augmented |

The analyzed parameters were: detection of the PCN mRNA or protein; effects on the *in vitro* yeast growth; effects on the transition from yeast to mycelium; sensitivity to killing by macrophages; effects on the infection pathogenesis, and grade of virulence. The virulence was inferred from the set of analyzed parameters.

The observation that fungal growth was not affected by overexpression of PCN shows that the activity of this enzyme is needed for normal growth [22], but its increase has no surplus effect. As such, our data further suggests that the PCN chitinase activity is required for chitin hydrolyses allowing the separation of the budding of the daughter cells.

We found that overexpression of PCN had an acceleration effect on the transition from yeasts to hypha, compared to wt-PCN yeasts. Accordingly, PCN-knocked-down yeasts underwent transition blockage [18]. Delayed inhibition of yeast-to-hyphae transition was previously associated to other *P. brasiliensis*-components silencing, such as PbSconC [16], and Pb14-3-3 [28].

Our current results could no be in line with the previous demonstrations that administration of exogenous rPCN to mice confers resistance against PCM [44, 45], if one could take only the presence of PCN, and not account for the fungal-biological function of this protein. Indeed, increased endogenous PCN, to which ov-PCN yeast-infected mice were exposed, brought no beneficial effect to the host; on the contrary, animals were severely ill, with high fungal burden, serious pulmonary lesions, and high mortality score (see model shown in Fig 6). It seems that the endogenous PCN release is not sufficient to promote an immunomodulation that triggers protection, however the overexpression leads to alteration in the chitin processment profile of the mutant strains. Involvement of PCN chitinase activity has been reported; indeed, we verified the involvement of the substrate of PCN chitinase, which is chitin of the yeast cell wall, and the fragments generated by chitin hydrolysis. Several studies, published in the last decade, have explored particles derived from chitin hydrolysis concerning their property of stimulating macrophages to produce cytokines. The release of IL-10 and TNF-α was regularly examined as markers of the anti-inflammatory or inflammatory microenvironment, generated by the stimulation of macrophages by chitin particles. The response variability is dependent on the size and concentration of the stimulating chitin particles, as well as on the innate immunity receptor(s) targeted by the chitin particles. It is well known that an anti-inflammatory microenvironment, marked by a high IL-10 detection [36], promotes the development of non-protective immune response, which results in severe pulmonary damage in *P. brasiliensis*-infected hosts. A study conducted by Cunha, et al. showed that PBMCs with GG genotype secrete smaller amounts of TNF-α than carriers of AA or GA genotype after infection with *A. fumigatus*. In this sense, GG homozygotes generate fewer inflammatory responses. The dichotomy between IL-10 and TNF-α production according to genotypes rs1800896 was confirmed in human macrophages with the same genotype changes and, extended to other pro-inflammatory cytokines, such as IL-6, IL-1β, and IL-8 [49]. In our studies, chitin particles obtained from the supernatant of ov-PCN yeasts were the smallest measured fragments and induced the production of IL-10 when stimulating macrophages. Otherwise, chitin particles derived from the wt-PCN yeasts had a broader size distribution and had stimulated macrophages to a prominent TNF-α and IL-1β production. The availability of high PCN amounts, as occurs in ov-PCN yeasts, leads to a more effective chitin cleavage and stimulation of IL-10 release by macrophages. This justifies the severity of infection with ov-PCN *P. brasiliensis* yeasts and the accompanying pulmonary damage compared to those caused by wt-PCN yeasts.

**Fig 6.**
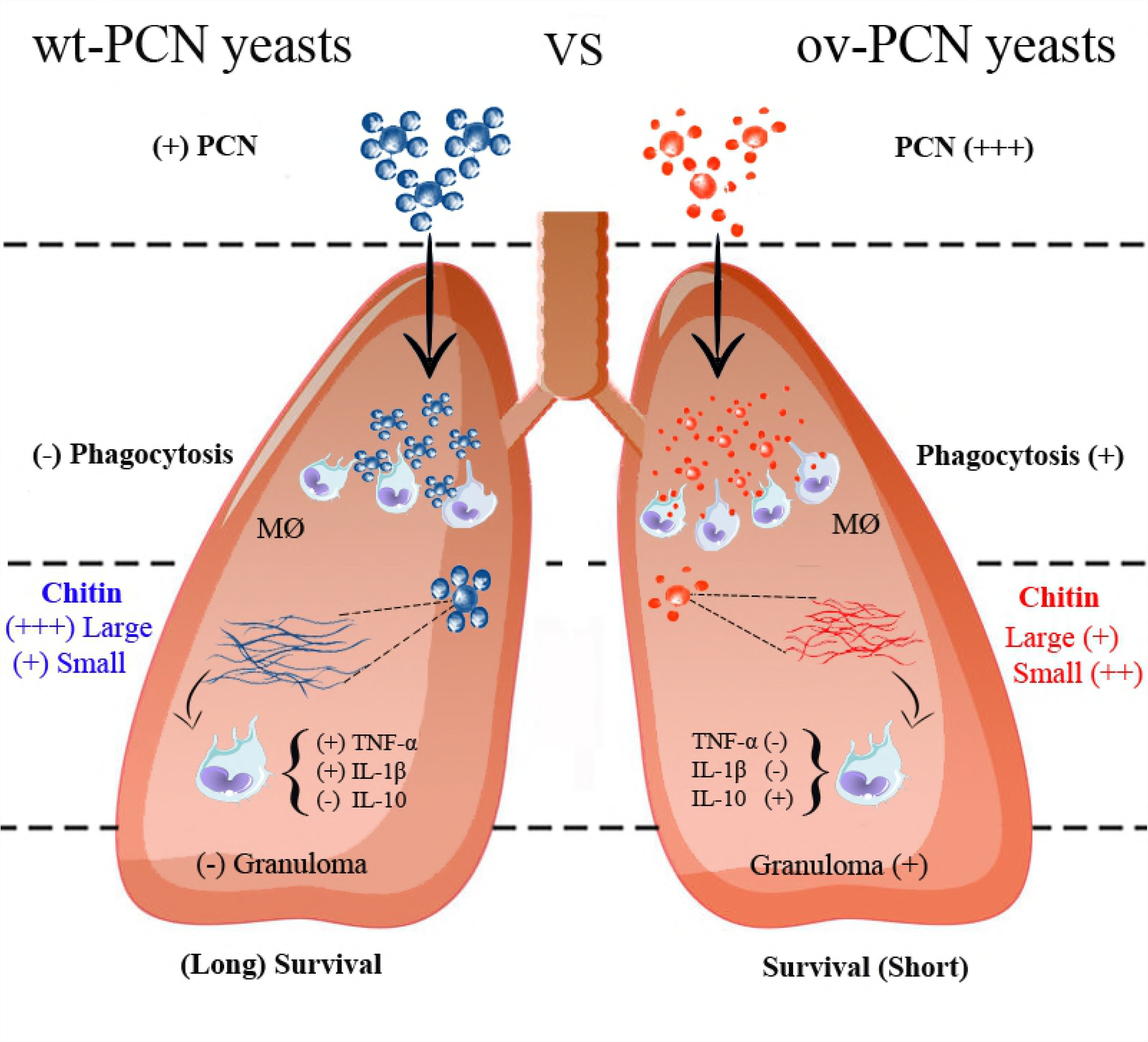
Paracoccin (PCN) acts as a virulence factor of *P. brasiliensis*, as demonstrated by comparing the performance of ov-PCN and wt-PCN yeasts in *in vitro* and *in vivo* assays. The study contributes to an updated understanding of the role of PCN in *P. brasiliensis* infection. PCN confers yeast resistance to the macrophage fungicidal mechanisms, propitiates chitinase activity, promotes an efficient hydrolysis of chitin of the yeast cell wall and generates very small chitin particles. Once released in the extracellular milieu, the small particles activate macrophages to produce anti-inflammatory cytokines. The generated microenvironment induces a non-protective immune response against the fungus. Such a chain of events is responsible, at least in part, for the severe pulmonary disease produced following the mice infection with *P. brasiliensis* yeasts overexpressing Paracoccin.

In summary, the severe pulmonary disease caused by ov-PCN yeasts is mechanistically related to the anti-inflammatory activity of macrophages toward the chitin with an higher level of hydrolysis and the consequent no immune response against the fungus (see model in Fig 6). Our data demonstrate that the macrophage cytokines-response depends also on the concentration of chitin particles used to stimulate macrophages. When the particles were used at 25 to 100 µg/mL, there was a better discrimination between the levels of IL-10 and TNF-α produced by macrophages stimulated by chitin particles derived from ov-PCN and wt-PCN yeasts. This observation is consistent with that of Wagener et al. (2014), when working with chitin particles derived from *C. albicans* cell wall. Besides the mechanism involving macrophage activation by chitin particles, a higher number of ov-PCN yeasts was phagocytized by macrophages than that of wt-PCN yeasts, but the cell fungicidal activity was not effective. In conclusion, by overexpressing PCN in yeasts, we demonstrated that PCN is a virulence factor of *P. brasiliensis*, able to contribute a severe pulmonary disease. The mechanisms involved in the severe course of murine infection with the PCN-overexpressing yeasts are linked to its ability to hydrolyze cell wall chitin to very small particles. They induce macrophages to generate an anti-inflammatory environment, able to attenuate the immune response, thereby reducing the ability of the host to combat the fungal pathogen (see model in Fig 6). Further investigation is required for a complete understanding of the role of PCN in *P. brasiliensis* infection.

### A model for the mechanism by which PCN acts as a virulence factor of *P. brasiliensis* yeasts

## MATERIALS AND METHODS

### Ethics statement

All experiments were conducted in accordance to the Brazilian Federal Law 11,794/2008 establishing procedures for the scientific use of animals, and State Law establishing the Animal Protection Code of the State of Sao Paulo. All efforts were made to minimize suffering, and the animal experiments were approved by the Ethics Committee on Animal Experimentation (Comissão de Ética em Experimentação Animal – CETEA) of the Ribeirao Preto Medical School, University of Sao Paulo (protocol number 061/2016), following the guidelines of the National Council for Control of Animal Experimentation (Conselho Nacional de Controle de Experimentação Animal – CONCEA).

### Western blot

The *P. brasiliensis* wt-PCN yeasts were grown in BHI broth at 37 °C for 72 h (exponential phase). Then, the wt-PCN yeasts were co-cultured with the human monocytes THP-1 cell line obtained from the American Type Culture Collection [ATCC] during 24 h. The western blot analyses were performed by the use of 12 % SDS-PAGE to separate the soluble elements in supernatants of co-cultures. The components were ran electrophoretically and the proteins were electrotransferred to a polyvinylidene difluoride membrane (Hybond-P™, Amersham GE Healthcare) at 50 V and a current of 90 mA overnight using a transfer buffer (1.9% Tris base, 9.1% glycine). The non-specific interactions were blocked by incubating the membranes with TBS-T 1× (20 mM Tris-HCl, 150 mM NaCl, 0.05% Tween 20 [pH 7.6]) containing 5% skim milk powder (Molico^®^) for 1 h under slow stirring at room temperature. Subsequently, the membrane was incubated with anti-PCN IgY polyclonal antibody diluted 1:3,000 in TBS-T. The membrane was washed five times with TBS-T and incubated for 1 h at room temperature with the anti-IgY secondary antibody conjugated to peroxidase (Sigma-Aldrich^®^, St. Louis, MO, USA), diluted 1:1,000. After five washes with TBS-T were performed, the membrane was immersed in a fresh mixture of the DAB peroxidase substrate kit SK4100 (Vector Laboratories, Burlingame, CA, USA). Distilled water was used to stop the reaction. The images were analyzed using the chemiDocTMMP Imaging System (Bio-Rad, USA).

### Construction of ov-PCN cassettes

Plasmids and genomic DNA extraction, recombinant DNA manipulations, and *E. coli* transformation procedures were performed as described elsewhere [23]. DNA from the *P. brasiliensis* strain Pb18 was extracted from yeast cultures during exponential growth, and a high-fidelity proofreading DNA polymerase (NZYTech, Portugal) was employed to amplify 1025 bp with exon and intron sequences corresponding to the PADG_03347 <u>(Gene ID: 22582669</u>) sequence. The primer sequences were synthesized by Sigma-Aldrich, and the sequences are gParacoccin forward (5’-GGCGCGCCATGGCCTTCGAAAATCAG-3’) and gParacoccin reverse (5’-CTCGAGTTACCATGAACTCGTCGA-3’). The PCN cassette was inserted in a pCR35-RHO2 vector under control of the Rho (Ras homology) GTPase 2 (*RHO2*) promoter region from *P. brasiliensis*. The pCR35 plasmid was digested with the restriction enzymes *AscI* and *XhoI*. The amplified *P. brasiliensis* PADG_03347 sequences were digested with the same enzymes and cloned into Asc*I-XhoI-*digested pCR35. Genomic PCN expression cassettes were amplified with the primers gParacoccin forward and reverse and digested with the *KpnI* restriction enzyme. Subsequently, they were cloned into the t-DNA region of the binary vector pUR5750, previously digested with *KpnI*. Resulting vectors were introduced into A. *tumefaciens* LBA1100 ultracompetent cells by electroporation, as previously described [23]. Transformants were isolated by selection on kanamycin at 100 μg/mL.

### ATMT of *P. brasiliensis*

Insertion of recombinant T-DNA harboring the ov-PCN cassettes and a hygromycin B resistance marker into the genome of *P. brasiliensis* yeast cells was accomplished by ATMT [23, 46]. Briefly, *A. tumefaciens* strains carrying the binary vector pUR5750 harboring the ov-PCN cassettes were grown in Luria Bertani (LB) broth containing kanamycin (100 µg/mL), rifampicin (20 µg/mL), and spectinomycin (250 µg/mL) at 29 °C with aeration in a mechanical shaker at 150 rpm overnight. Bacterial cells were spun-down, washed with induction medium (IM) [23, 46] and resuspended in 10 mL of IM with 0.2 M acetosyringone (AS) (Sigma-Aldrich) and antibiotics. Cells were grown until they provided an optical density (OD_600_) of approximately 0.8. Then, yeast cells were grown in BHI broth and harvested during the exponential growth phase. The cells were washed with IM and adjusted to a final concentration of 1×10^8^ cells/mL, estimated by using direct microscopic counts in a Neubauer chamber. For co-cultivation, 1:1 and 10:1 ratios of *A. tumefaciens* and *P. brasiliensis* cells were spotted on sterile Hybond N filters (Amersham Biosciences, USA), in solid IM with AS and antibiotics, dried in a safety cabinet for 30 min in the dark and incubated for 3 days at 25 °C. Membranes were then transferred to BHI broth containing cefotaxime (200 µg/mL), and cells were dislodged with the aid of a spatula before incubated for 48 h at 37 °C in a shaker (220 rpm). *P. brasiliensis* cells were then plated on BHI broth, supplemented with the appropriate antibiotic (hygromycin 75 µg/mL), and grown at 37 °C for 15 days. The obtained transformed yeasts were kept for further analysis. We used *P. brasiliensis* yeasts transformed with an EV, pUR5750, as a control in assays carried out in this study.

### Microorganisms and culture medium

All strains used in this study are listed in Table 1.

**Table 1.** Strains and plasmids used in this study.

| Strains | Genotype | Source |
| --- | --- | --- |
| *A. tumefaciens*<br>LBA1100 | C58 chromosomal background<br>Rifampicin<sup>R</sup>,<br>pectinomycin<sup>R</sup>; pAL1100<br>or pTiB6 ΔTL, ΔTR,<br>Δtra, Δocc | [50] |
| *E. coli* DH5α | F<sup>-</sup> endA1 glnV44 thi-1<br>recA1 relA1 gyrA96<br>deoRnupGΦ80dlacZΔM<br>15 Δ(*lacZYA-argF*)<br>U169, hsdR17(rK-<br>mK+), λ- | [51] |
| *P. brasiliensis* | Pb18 | USP-RP, Brazil |
| Plasmids | Selection/genetic<br>Markers |  |
| pUR5750 | Hygromycin<sup>R</sup> | [52] |
| pCR35::P<sub>RHO2</sub>-gPCN | Kanamycin<sup>R</sup> | This study |
| pUR5750::P<sub>RHO2</sub>-gPCN | Hygromycin<sup>R</sup> | This study |

*P. brasiliensis* yeast cells were maintained at 36 °C by periodic subculturing on BHI medium (Duchefa, Netherlands), supplemented with 1% glucose and 1.5% v/v agar or Dulbecco’s modified Eagle’s medium (DMEM) (Sigma-Aldrich). For the assays performed in this study, yeast cells were grown in liquid BHI broth at 36 °C with aeration on a mechanical shaker at 200 rpm. Their viability was evaluated by fluorescein diacetate and ethidium bromide staining [53] and only yeasts suspensions with viability higher than 90% were included in the study. The LBA1100 strain of *A. tumefaciens* was used as the recipient of the binary vectors herein constructed [25]. Bacterial cells were maintained at 28 °C in LB broth (Kasvi, Italy), containing kanamycin (50 µg/mL), spectinomycin (250 µg/mL) and rifampicin (20 µg/mL). *Escherichia coli DH5α* competent cells, grown at 37 °C in LB broth supplemented with antibiotics (kanamycin 50 µg/mL; ampicillin 100 µg/mL), were used as hosts for plasmid amplification and cloning.

### Gene and ov-PCN analysis

Yeast cells of *P. brasiliensis* obtained from a single colony were inoculated and cultured in BHI broth at 37 °C (200 rpm), until the exponential growth phase. The culture medium was refreshed once, after 4 days. Total RNA obtained from ov-PCN and wt-PCN yeasts and isolated according to the TRIzol protocol (NZYTech, Portugal). The RNA samples were subsequently incubated for 20 min at 37 °C with 2 U of DNase I (Roche, Germany). The absence of DNA contamination in the samples was checked by the lack of conventional PCR amplification of the *GP43* gene in the isolated RNA. Total RNA (1 μg) was reversely transcribed using the NZYTech Reverse Transcriptase cDNA Synthesis kit (NZYTech, Portugal) according to the manufacturer’s instructions. As a template for real-time quantification, cDNA (1 µL) of was used in the SoFast EvaGreen SuperMix (Bio-Rad) according to the manufacturer’s instructions. Quantitative Real-time PCR was carried out on a CFX96 Real-Time System (Bio-Rad), using threshold cycle (Ct) values for β-tubulin (*TUB2)* and ribosomal protein (*L34*) transcripts as endogenous references. The primer sequences were synthesized by NZYTech (Portugal) and the sequences are β-tubulin forward (ACGCTTGCGTCGGAACATAG), β-tubulin reverse (ACCTCCATCCAGGAACTCTTCA), L34 forward (CGGCAACCTCAGATACCTTC) and L34 reverse (GGAGCCTGGGAGTATTCACG). mRNA differential ov-PCN was evaluated by normalizing gPCN Ct values with the reference and comparing the ratio among the tested samples. All the measurements were performed in triplicate.

### Molecular detection of the hygromycin resistance gene (hph)

Genomic DNA from ov-PCN yeasts and control yeast cells were isolated according to the glass beads protocol described by Van Burik [54]. In order to confirm the presence of the hygromycin B resistance cassette, PCR analysis was carried out to detect an HPH amplification product (1000 bp) using the hph forward (AACTCACCGCGACGTCTGTCGA) and hph reverse (CTACACAGCCATCGGTCCAGA) primers (data not shown). PCR amplification included 30 cycles of 5 min at 94 °C, for denaturing, 40 s at 55 °C for annealing, and 1 min 10 s at 72 °C for extension. The reaction products were analyzed on 1% agarose gel and visualized with SYBR Green, under UV light.

### Enzymatic activity of ov-PCN yeasts

The NAGase activity of ov-PCN yeasts was assayed as previously described [20, 55]. Briefly, the substrate *p*-nitrophenyl-N-acetyl-β-D-glucosaminidase (100 µL, 5 mM; Sigma-Aldrich) was mixed with of 0.1 M sodium acetate, pH 5.5 (350 µL) and a sample of the supernatant cultures (50 µL). As a negative control, we used the vehicle medium. The reaction was incubated for 16–18 h at 37 °C, in medium to which we have added 0.5 M sodium carbonate (1 mL). The enzyme activity values were estimated by using a microplate reader at 405 nm (Power Wave X, BioTek Instruments, Inc.). The NAGase activity in the supernatant of wt and transformed yeast cultures was considered to indicate the relative concentration of PCN produced by each *P. brasiliensis* strain.

### Growth curve and viability assay

Growth curves were performed in DMEM (10 mL) by inoculating, at 48 h, washed fungal cells (1×10^6^); the OD_600nm_ was adjusted to reach the value 0.2. Cellular density was measured in triplicates in a spectrophotometer (Ultrospec^®^3000pro, GE Healthcare), at 24, 48, 72, 96, 120, 144, 168, 192, 216 and 240 h of growth, for wt-PCN and ov-PCN yeasts of the strain Pb18, as previously described [26]. The viability of wt-PCN and ov-PCN yeasts was determined by fluorescein diacetate-ethidium bromide staining [53] and detected by fluorescence microscopy (excitation bands: blue-BP480/40; green-BP515-560) (DMI6000B, Leica). Only suspensions containing at least 90% of viable yeasts were included in the studies. The experiments were performed in triplicate.

### Fluorescence labeling confocal microscopy

Confocal microscopy was performed on yeasts that were harvested from wt-PCN, EV, and ov-PCN yeasts, grown in BHI (Kasvi) at 37 °C for 72 h (exponential phase) and separated by centrifugation at 4,000×*g* at 25 °C for 6 min. Yeast cells were fixed using 3.7% para-formaldehyde in phosphate-buffered saline (PBS) pH 7.2 at 25 °C. After 1 h, yeasts were washed with PBS containing 1% glycine. For labeling, blocked samples with 1 mL PBS containing 1% BSA, at 25 °C for 1 h, were incubated with the biotin-conjugated anti-PCN IgY antibody (1:50) for 1 h at 25°C. Then, the yeast cells were incubated with the streptavidin Alexa 594-conjugated antibody (Thermo Fisher Scientific), for 1 h at 25 °C. The cell images were acquired in the LSM 780 AxioObserve Inverted Microscope (Carl Zeiss, Jena, Germany), available in a multi-user institutional facility, installed in the Cell and Molecular Biology Department. For all samples obtained from wt-PCN, EV and ov-PCN yeasts, identical photomultiplier gain and laser power were employed. Images were recorded and analyzed offline with the aid of the software ImageJ [(W. S. Rasbanda/National Institute of Health, Bethesda, MD, USA (http://rsb.info.nih.gov./iji/)].

### Infection induction in mice

We performed the infection through intranasal or intravenous inoculation of yeast cells, collected from exponentially growing batch cultures in BHI broth, and counted in a Neubauer chamber. Animals of each group (n=5) were inoculated with 2×10^6^ yeast cells from the wt-PCN or ov-PCN *P. brasiliensis* strains, contained in 40 µL PBS, as specified in Table 2.

**Table 2.**
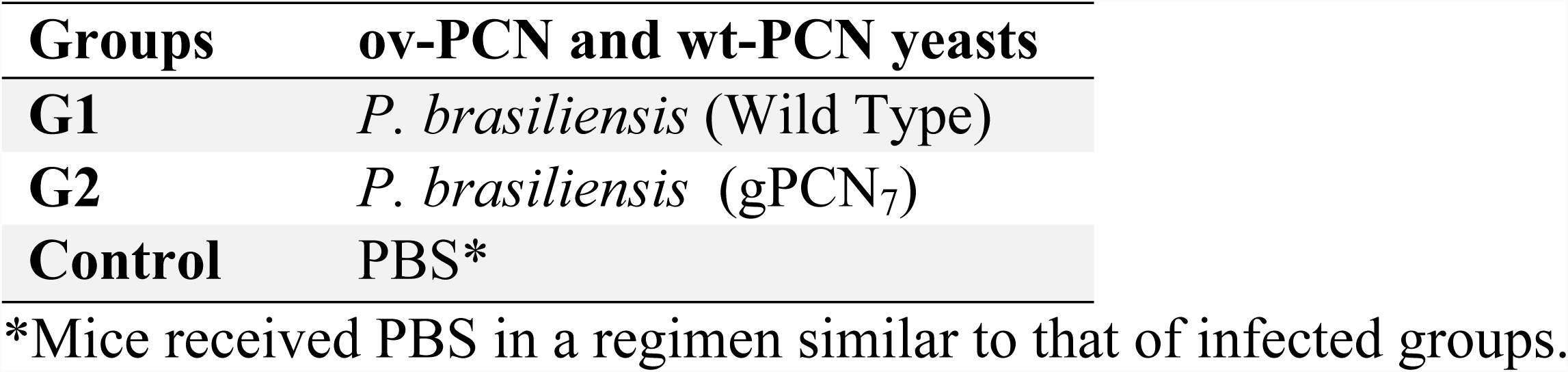
ov-PCN groups intranasal infection.

Uninfected control mice were inoculated with vehicle alone, under the same conditions as the infected group. At 30 days post-infection, mice were euthanized, and their lungs were harvested for analysis. For the survival studies, each mouse of a group (n=10) was intravenously inoculated with 1×10^6^ yeast cells of wt-PCN or ov-PCN strain, contained in 15 µL PBS (Table 3). Mice were monitored daily for mortality during 70 days post-infection.

**Table 3.** ov-PCN groups intravenously infection.

| Groups | ov-PCN and wt-PCN yeasts |
| --- | --- |
| G1 | *P. brasiliensis* (Wild Type) |
| G2 | *P. brasiliensis* (gPCN<sub>7</sub>) |

### Pulmonary fungal burden in ov-PCN infected mice

The pulmonary fungal burden was evaluated by counting the number of CFU recovered from a similar lung fragment of each infected mouse. The fragment was aseptically removed, weighed, and homogenized in 1.0 mL sterile PBS in a tissue homogenizer (Ultra-Turrax T25 Basic; IKA Works, Inc., Wilmington, USA). The supernatant was plated on solid BHI medium supplemented with 4% (v/v) heat-inactivated fetal bovine serum (Invitrogen, Life Technologies, Camarillo, CA, USA) and gentamicin at 96 µg/mL (Gibco, Grand Island, USA). The plates were incubated at 36 °C for 7 days, after which the colonies were counted.

### Lung histopathological analysis

An excised lung of each mouse was fixed in 10% para-formaldehyde for 24 h, dehydrated in ethanol, diaphanized in xylene, and embedded in paraffin. Histological sections of 6 μm thickness were hematoxylin-eosin stained for histological analysis. Images were acquired in a ScanScope multi-user scanner Lab, using an optical microscope (Olympus VS120 in a BX 61 microscope, Axiophot Photo microscope; Carl Zeiss GmbH, Germany) and a camera (JVC TK-1270; Victor Company of Japan Ltd., Japan).

### Fungal phagocytosis and killing by murine macrophages

Yeast cells from ov-PCN and wt-PCN strains, grown in BHI broth at 37 °C for 72 h (exponential phase), were co-cultured with RAW 264.7 murine macrophages (obtained from the American Type Culture Collection [ATCC]) were cultivated in complete RPMI (Sigma-Aldrich), supplemented with 10% fetal bovine serum (FBS), 2 mM L-glutamine and 100 µg/mL streptomycin/ampicillin. The fungal inoculum was prepared as described for the previous item.

To assess their phagocytosis, the macrophages were distributed (5×10^5^ cells/well) in a 24-well microplate (Costar, Corning Inc., Corning, NY, USA) and incubated with yeast cells (5×10^4^ yeasts/well) for 4 h at 37 °C (yeast-to-macrophage ratio of 1:10) in a 5% CO_2_ atmosphere, and the co-culture supernatant was discarded. The macrophage monolayers were lysed with ice-cold water, and the cell lysate was plated on BHI agar broth supplemented with 1% glucose, for 7 days at 37 °C.

To estimate the macrophage fungicidal activity, viable yeast cells were co-cultured with macrophages at 37 °C in a 5% CO_2_ atmosphere for 4 h. The culture supernatant was discarded, and the cells were gently rinsed with PBS and incubated in RPMI (Sigma-Aldrich) supplemented with 10% FBS (HyClone) at 37 °C in a 5% CO_2_ atmosphere. After 48 h of incubation, the culture supernatant was discarded, macrophages were lysed using ice-cold water, and the cell lysate was serially diluted and cultured, for 7 days at 37 °C, in solid BHI broth supplemented with 1% glucose. The analysis was performed by counting the number of CFU.

### Chitin capture from supernatants of yeast cultures

WGA (Sigma-Aldrich) was coupled to CNBr-Activated Sepharose 4B beads (GE Healthcare), according to the manufacturer’s instructions. The conjugation yield was about 90% (4.5 mg protein per mL of resin). The lectin-resin coupling was stabilized through crosslinking with glutaraldehyde [56]. Two columns of 1-mL resin were packaged, each one dedicated to either chitin derived from wt-PCN or transformed yeasts. The yeast cultures (ov-PCN or wt-PCN yeasts) were grown in BHI broth, for 7 days, at 37 °C and 180 rpm. After centrifugation of the cultures at 3300*×g* for 10 min, at room temperature, the cultures’ supernatants were loaded (10 mL) in the columns. After incubation for 90 min, under slow rotation at room temperature, the unbound material was eluted with PBS, while the bound material was subsequently eluted with 0.5 M NaCl in PBS. The PBS-eluate material was centrifuged and re-chromatographed (same protocol) to overcome an eventual overloading of the column in the first procedure. The NaCl 0.5 M-eluate (WGA-bound material) was ultradiafiltered against water (OS20LX) in a 0.05-µm VM membrane (Millipore^®^). The resultant preparations derived from wt-PCN and ov-PCN yeasts were stored at 4 °C. They were named wt-PCN chitin and ov-PCN chitin.

### Carbohydrate quantification

The samples’ carbohydrate content was determined through the modified Dubois Colorimetric Method [57], which was especially useful to analyze the chitin purified preparations. The standard curve concentration was constructed by serially diluting in water a starting solution of glucose (5 mg/mL). The reaction was performed in 2 mL microtubes by using 500 µL samples, concentrated sulfuric acid (800 µL) and an 80% phenol solution (50 µL). Optical density (490 nm) was read in a Power Wave X microplate scanning spectrophotometer (BioTek Instruments, Inc.). All reactions were done in triplicates. The sugar concentration was expressed as micrograms.

### Size of chitin particles and thickness of cell wall determined by transmission microscopy

The column containing immobilized WGA was used for purification of chitin particles from ov-PCN and wt-PCN yeasts (fixed in glutaraldehyde 2% and para-formaldehyde 4% and prepared on SPURR resin as according to the manufacturer’s instructions). Chitin particles and yeasts were analyzed by transmission electron microscopy (Jeol JEM-100 CXII equipped with Hamamatsu digital camera ORCA-HR; magnification 20,000, 80,000, and 200,000 ×). The images were analyzed in ImageJ software (Image-adjust-threshold-(select the area) -analyze particles) and the thickness of the cell wall in the ov-PCN and wt-PCN yeasts was determined.

### Activation of BMDMs by chitin particles

BMDMs were prepared from femurs and tibias of BALB/c mice after flushing with RPMI medium to release the bone marrow cells. These cells were cultured in a 100 mm × 20 mm suspension culture dish (Corning) with 10 mL of RPMI 1640 broth supplemented with supernatant from L929 cell cultures (20% FCS and 30% supernatant from L929 cell cultures, LCCM, L929-cell conditioned medium). In the fourth day, 10 mL of RPMI medium supplemented with supernatant purified from L929 cell cultures were added. In the seventh day, the non-adherent cells were removed, and the dishes were washed with PBS. After that, the adherent cells (macrophages) were removed by heat shock with cold PBS (for 10 min) and washed twice with PBS. The concentration of the cells was determined in a Neubauer chamber, and the BMDMs (1×106/mL) were cultured in 96-well microplates for 24, 48 and 72 h at 37 °C in the presence with a pool of particles (2.5, 7.5, 25, 50, and 100 µg/mL) or medium alone. The supernatant was used for quantification of cytokines.

### Quantification of cytokines

Supernatants of stimulated of BMDMs macrophages were assessed for their levels of TNF-α, IL-1β, and IL-10. The cytokines were detected by an enzyme-linked immune sorbent assay (ELISA) using an OptEIA kit (Pharmingen, SanDiego, CA, USA), according to the manufacturer’s instructions. Standard curves allowed determining cytokine concentrations in pg/mL. The absorbance was read at 450 nm using the Power Wave X microplate scanning spectrophotometer (BioTek Instruments, Inc.).

### Statistical analysis

All statistical analysis was performed by using the GraphPad Prism Software version 6.0. Data are reported as the mean ± standard error of the mean (SEM). All the results of *in vivo* and *in vitro* assays are representative of three independents assays. Five or ten animals constituted each control or experimental group. For all data analysis, statistical significance was considered at the level of followed by One-Way ANOVA test or Mann-Whitney test. Differences with \*\*\*\**p*<0.0001, \*\*\**p*<0.001, \*\**p*<0.01 and \**p*<0.1 were considered statistically significant.

## ACKNOWLEDGEMENTS

RAG was supported by FAPESP (Project number: 2014/22561-7) and CAPES (Project number: 99999.010699/2014-07). FR, AC and CC were supported by the Northern Portugal Regional Operational Programme (NORTE 2020), under the Portugal 2020 Partnership Agreement, through the European Regional Development Fund (FEDER) (NORTE-01-0145-FEDER-000013), and the Fundação para a Ciência e Tecnologia (FCT) (SFRH/BPD/96176/2013 to CC and IF/00735/2014 to AC). We are grateful to Mrs Vani M. Alvez, Roberta Rosales, Elizabet Rosa, Maria Dolores Ferreira, Patricia V. Bonini, and Mr. José Maulin for helpful intellectual discussions.

## SUPPORTING INFORMATION

**S1 Fig.**
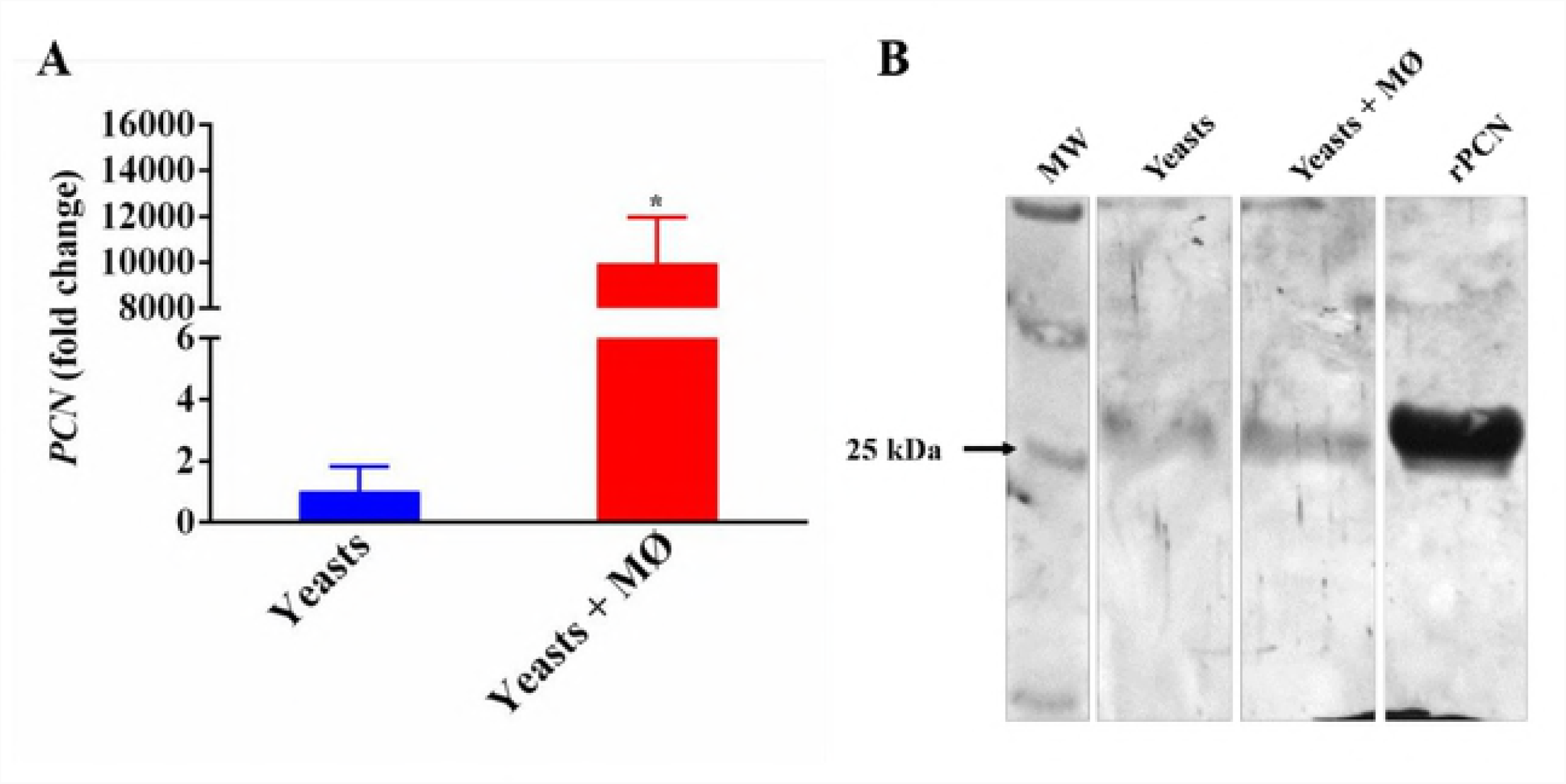
Chitin isolated from supernatants of ov-PCN and wt-PCN yeasts induces macrophages to release cytokines in a dose-dependent manner. BMDMs (1×10^6^ cells/mL) were stimulated for 24, and 72 h with chitin particles captured from the supernatants of yeast cultures (ov-PCN or wt-PCN yeasts). The macrophages supernatants were assessed by ELISA for cytokine levels (pg/mL) by dose-response of chitin concentration (2.5 µg/mL to 100 µg/mL). **(A and D)** TNF-α. **(B and E)** IL-1β, and **(C and F)** IL-10. The supernatants of unstimulated macrophages or LPS-stimulated macrophages (1 mg/mL) were used as negative and positive controls, respectively. The results are expressed as mean ± SEM and were compared to the values obtained by the negative control (cells) through one-way analysis of variance, followed by One-Way ANOVA test. \*\*\*\**p*<0.0001, \*\*\**p*<0.001, \*\**p*<0.01 and \**p*<0.1.

## REFERENCES

1. San-Blas, G., G. Niño-Vega, and T. Iturriaga, Paracoccidioides brasiliensis and paracoccidioidomycosis: Molecular approaches to morphogenesis, diagnosis, epidemiology, taxonomy and genetics. Medical Mycology, 2002. 40(3): p. 225–242.

2. Rappleye, C.A. and W.E. Goldman, Defining virulence genes in the dimorphic fungi. Annu Rev Microbiol, 2006. 60: p. 281–303.

3. Mendes, J.F., et al., Paracoccidioidomycosis infection in domestic and wild mammals by Paracoccidioides lutzii. Mycoses, 2017. 60(6): p. 402–406.

4. Teixeira, M.M., et al., Phylogenetic analysis reveals a high level of speciation in the Paracoccidioides genus. Mol Phylogenet Evol, 2009. 52(2): p. 273–83.

5. Theodoro, R.C., et al., Genus paracoccidioides: Species recognition and biogeographic aspects. PLoS One, 2012. 7(5): p. 30.

6. Puccia, R., et al., Exocellular components of Paracoccidioides brasiliensis: identification of a specific antigen. Infect Immun, 1986. 53(1): p. 199–206.

7. De Camargo, Z., et al., Production of Paracoccidioides brasiliensis exoantigens for immunodiffusion tests. J Clin Microbiol, 1988. 26(10): p. 2147–51.

8. Giannini, M.J., et al., Antibody response to the 43 kDa glycoprotein of Paracoccidioides brasiliensis as a marker for the evaluation of patients under treatment. Am J Trop Med Hyg, 1990. 43(2): p. 200–6.

9. Travassos, L.R., et al., Biochemistry and molecular biology of the main diagnostic antigen of Paracoccidioides brasiliensis. Arch Med Res, 1995. 26(3): p. 297–304.

10. Cisalpino, P.S., et al., Cloning, characterization, and epitope expression of the major diagnostic antigen of Paracoccidioides brasiliensis. J Biol Chem, 1996. 271(8): p. 4553–60.

11. Izacc, S.M., et al., Molecular cloning, characterization and expression of the heat shock protein 60 gene from the human pathogenic fungus Paracoccidioides brasiliensis. Med Mycol, 2001. 39(5): p. 445–55.

12. Cunha, D.A., et al., Heterologous expression, purification, and immunological reactivity of a recombinant HSP60 from Paracoccidioides brasiliensis. Clin Diagn Lab Immunol, 2002. 9(2): p. 374–7.

13. Fernandes, F.F., et al., Detrimental Effect of Fungal 60-kDa Heat Shock Protein on Experimental Paracoccidioides brasiliensis Infection. PLoS One, 2016. 11(9).

14. Fernandes, V.C., et al., Protective effect of rPb40 as an adjuvant for chemotherapy in experimental paracoccidioidomycosis. Mycopathologia, 2012. 174(2): p. 93–105.

15. Fernandes, V.C., et al., Additive effect of rPb27 immunization and chemotherapy in experimental paracoccidioidomycosis. PLoS One, 2011. 6(3): p. 0017885.

16. Menino, J.F., et al., P. brasiliensis virulence is affected by SconC, the negative regulator of inorganic sulfur assimilation. PLoS One, 2013. 8(9).

17. Almeida, A.J., et al., Cdc42p controls yeast-cell shape and virulence of Paracoccidioides brasiliensis. Fungal Genet Biol, 2009. 46(12): p. 919–26.

18. Fernandes, F.F., et al., Impact of Paracoccin Gene Silencing on Paracoccidioides brasiliensis Virulence. MBio, 2017. 8(4): p. 00537–17.

19. Coltri, K.C., et al., Paracoccin, a GlcNAc-binding lectin from Paracoccidioides brasiliensis, binds to laminin and induces TNF-alpha production by macrophages. Microbes Infect, 2006. 8(3): p. 704–13.

20. dos Reis Almeida, F.B., et al., Paracoccin from Paracoccidioides brasiliensis; purification through affinity with chitin and identification of N-acetyl-beta-D-glucosaminidase activity. Yeast, 2010. 27(2): p. 67–76.

21. Dos Reis Almeida, F.B., et al., Influence of N-glycosylation on the morphogenesis and growth of Paracoccidioides brasiliensis and on the biological activities of yeast proteins. PLoS One, 2011. 6(12): p. 21.

22. Ganiko, L., et al., Paracoccin, an N-acetyl-glucosamine-binding lectin of Paracoccidioides brasiliensis, is involved in fungal growth. Microbes Infect, 2007. 9(6): p. 695–703.

23. Almeida, A.J., et al., Towards a molecular genetic system for the pathogenic fungus Paracoccidioides brasiliensis. Fungal Genet Biol, 2007. 44(12): p. 1387–98.

24. Hernandez, O., et al., A 32-kilodalton hydrolase plays an important role in Paracoccidioides brasiliensis adherence to host cells and influences pathogenicity. Infect Immun, 2010. 78(12): p. 5280–6.

25. Menino, J.F., A.J. Almeida, and F. Rodrigues, Gene knockdown in Paracoccidioides brasiliensis using antisense RNA. Methods Mol Biol, 2012. 845: p. 187–98.

26. Torres, I., et al., Inhibition of PbGP43 expression may suggest that gp43 is a virulence factor in Paracoccidioides brasiliensis. PLoS One, 2013. 8(7).

27. Torres, I., et al., Paracoccidioides brasiliensis PbP27 gene: knockdown procedures and functional characterization. FEMS Yeast Res, 2014. 14(2): p. 270–80.

28. Marcos, C.M., et al., Decreased expression of 14-3-3 in Paracoccidioides brasiliensis confirms its involvement in fungal pathogenesis. Virulence, 2016. 7(2): p. 72–84.

29. Tamayo, D., et al., Identification and Analysis of the Role of Superoxide Dismutases Isoforms in the Pathogenesis of Paracoccidioides spp. PLoS Negl Trop Dis, 2016. 10(3).

30. Desjardins, C.A., et al., Comparative genomic analysis of human fungal pathogens causing paracoccidioidomycosis. PLoS Genet, 2011. 7(10): p. 27.

31. Munoz, J.F., et al., Genome update of the dimorphic human pathogenic fungi causing paracoccidioidomycosis. PLoS Negl Trop Dis, 2014. 8(12).

32. Oliveira, A.F., et al., Paracoccin distribution supports its role in Paracoccidioides brasiliensis growth and dimorphic transformation. PLoS One, 2017. 12(8).

33. Cano, L.E., et al., Pulmonary paracoccidioidomycosis in resistant and susceptible mice: relationship among progression of infection, bronchoalveolar cell activation, cellular immune response, and specific isotype patterns. Infect Immun, 1995. 63(5): p. 1777–83.

34. Itakura, Y., et al., Sugar-Binding Profiles of Chitin-Binding Lectins from the Hevein Family: A Comprehensive Study. Int J Mol Sci, 2017. 18(6).

35. Da Silva, C.A., et al., Chitin is a size-dependent regulator of macrophage TNF and IL-10 production. J Immunol, 2009. 182(6): p. 3573–82.

36. Da Silva, C.A., et al., TLR-2 and IL-17A in chitin-induced macrophage activation and acute inflammation. J Immunol, 2008. 181(6): p. 4279–86.

37. Becker, K.L., et al., Aspergillus Cell Wall Chitin Induces Anti- and Proinflammatory Cytokines in Human PBMCs via the Fc-gamma Receptor/Syk/PI3K Pathway. MBio, 2016. 7(3): p. 01823–15.

38. Dong, B., et al., Transformation of Fonsecaea pedrosoi into sclerotic cells links to the refractoriness of experimental chromoblastomycosis in BALB/c mice via a mechanism involving a chitin-induced impairment of IFN-gamma production. PLoS Negl Trop Dis, 2018. 12(2).

39. Dubey, L.K., et al., Induction of innate immunity by Aspergillus fumigatus cell wall polysaccharides is enhanced by the composite presentation of chitin and beta-glucan. Immunobiology, 2014. 219(3): p. 179–88.

40. Mora-Montes, H.M., et al., Recognition and blocking of innate immunity cells by Candida albicans chitin. Infect Immun, 2011. 79(5): p. 1961–70.

41. Wagener, J., et al., Fungal chitin dampens inflammation through IL-10 induction mediated by NOD2 and TLR9 activation. PLoS Pathog, 2014. 10(4).

42. Lee, C.G., et al., Chitin regulation of immune responses: an old molecule with new roles. Curr Opin Immunol, 2008. 20(6): p. 684–9.

43. Calich, V.L. and S.S. Kashino, Cytokines produced by susceptible and resistant mice in the course of Paracoccidioides brasiliensis infection. Braz J Med Biol Res, 1998. 31(5): p. 615–23.

44. Alegre, A.C., et al., Recombinant paracoccin reproduces the biological properties of the native protein and induces protective Th1 immunity against Paracoccidioides brasiliensis infection. PLoS Negl Trop Dis, 2014. 8(4).

45. Alegre-Maller, A.C., et al., Therapeutic administration of recombinant Paracoccin confers protection against paracoccidioides brasiliensis infection: involvement of TLRs. PLoS Negl Trop Dis, 2014. 8(12).

46. Michielse, C.B., et al., Agrobacterium-mediated transformation of the filamentous fungus Aspergillus awamori. Nat Protoc, 2008. 3(10): p. 1671–8.

47. Ricci-Azevedo, R., M.C. Roque-Barreira, and N.J. Gay, Targeting and Recognition of Toll-Like Receptors by Plant and Pathogen Lectins. Front Immunol, 2017. 8(1820).

48. Freitas, M.S., et al., Paracoccin Induces M1 Polarization of Macrophages via Interaction with TLR4. Frontiers in Microbiology, 2016. 7(1003).

49. Cunha, C., et al., IL-10 overexpression predisposes to invasive aspergillosis by suppressing antifungal immunity. J Allergy Clin Immunol, 2017. 140(3): p. 867–870.

50. Beijersbergen, A., et al., Conjugative Transfer by the Virulence System of Agrobacterium tumefaciens. Science, 1992. 256(5061): p. 1324–7.

51. Meselson, M. and R. Yuan, DNA restriction enzyme from E. coli. Nature, 1968. 217(5134): p. 1110–4.

52. de Groot, M.J., et al., Agrobacterium tumefaciens-mediated transformation of filamentous fungi. Nat Biotechnol, 1998. 16(9): p. 839–42.

53. Calich, V.L., A. Purchio, and C.R. Paula, A new fluorescent viability test for fungi cells. Mycopathologia, 1979. 66(3): p. 175–7.

54. van Burik, J.A., et al., Comparison of six extraction techniques for isolation of DNA from filamentous fungi. Med Mycol, 1998. 36(5): p. 299–303.

55. Almeida, F., et al., Toxoplasma gondii Chitinase Induces Macrophage Activation. PLoS One, 2015. 10(12).

56. Scher, M.G., W.G. Resneck, and R.J. Bloch, Stabilization of immobilized lectin columns by crosslinking with glutaraldehyde. Anal Biochem, 1989. 177(1): p. 168–71.

57. Dubois, M., et al., A colorimetric method for the determination of sugars. Nature, 1951. 168(4265): p. 167.

58. Nunes, L.R., et al., Transcriptome analysis of Paracoccidioides brasiliensis cells undergoing mycelium-to-yeast transition. Eukaryot Cell, 2005. 4(12): p. 2115–28.

